# Ubiquitin-Derived Diglycine Remnants as Chemical Handles for Chemoenzymatic Proteomics

**DOI:** 10.64898/2026.06.29.735432

**Authors:** Nicole R. Raniszweski, Myles I. Noel, Kamiel D. Beckley, Jordi C.J. Hintzen, George M. Burslem

## Abstract

Proteolytic processing is typically used to reveal post-translational modifications for mass spectrometric analysis. Here, we show that proteolysis of ubiquitinated proteins additionally generates a chemically addressable diglycine remnant that can be enzymatically remodeled. We exploit the ability of sortase to recognize ubiquitin-derived diglycine motifs as nucleophiles and ligate functionalized LPXTG peptides directly onto ubiquitin-remnant peptides. This reaction enables installation of a biotin affinity handle on defined ubiquitinated substrates and peptides derived from mammalian cell lysates, including lysine- and *N*-terminal ubiquitination. The resulting branched conjugates generate characteristic fragment ions that facilitate their identification. We further develop a bifunctional sortase donor containing an internal trypsin cleavage site, enabling affinity capture followed by conversion of enriched ubiquitin sites into a compact TGG mass-spectrometric signature. Alternatively, reversible sortase chemistry can regenerate the original diglycine-containing peptide. These studies establish ubiquitin-derived proteolytic remnants as latent chemical handles that can be converted into modular affinity and analytical tags, providing an antibody-independent approach to interrogating protein ubiquitination.

## Introduction

Post-translational modifications (PTMs) are an essential mechanism by which proteins in eukaryotes maintain controlled protein levels, adopt novel capabilities for interactions and signaling cascades, and perform countless other roles^1,2^. Among the suite of possible protein PTMs that can be added to a particular protein, ubiquitination has garnered significant attention, not least for its role in targeted protein degradation^3^. Ubiquitination involves the covalent attachment of ubiquitin (Ub), a stable 76-amino acid small protein, to proteins of interest. The process of ubiquitination is enacted by an enzyme cascade of E1, E2, and E3 proteins, and these ubiquitin marks are removed by deubiquitinases (DUBs)^4,5^. During the ubiquitination cascade, Ub is first activated in an ATP-dependent manner by the E1 before it is transferred to the protein of interest in an E2 and E3-dependent manner. Ubiquitination occurs predominantly on the sidechain of lysine residues forming an isopeptide linkage between Ub’s C-terminal glycine and the sidechain amine of the lysine; however ubiquitination can occur on the *N*-terminus of proteins, forming a peptide linkage^6,7^, as well as on non-lysine residues such as cysteine, serine, and threonine, thereby forming Ub thioester or esters, respectively^8–10^. Ub itself has 7 lysine residues that can act as ubiquitination sites to form polyubiquitin chains, thus adding an additional layer of complexity to the system. Despite over 50,000 protein ubiquitination sites having been identified in large proteomics screens^11–13^, our understanding of the scope of ubiquitination, as well as its roles in different biological contexts is far from complete; while many great strides have been made recently in available tools and reagents for detection of ubiquitination sites, there is still room for improvement when it comes to accurate and accessible means to interrogate protein ubiquitination.

The increased capability of mass spectrometry proteomic methods has revealed many aspects and targets of ubiquitination, with much of this data compiled in publicly available databases such as phosphositePlus^14^. Due to the low abundance of ubiquitinated proteins relative to the entire protein pool available in the cell, much of the ubiquitination data currently available was generated via ubiquitin-specific enrichment technologies, in which samples were treated to specifically select only for ubiquitinated proteins before profiling hits bearing the ubiquitin mark. Historically, this has been done in a variety of ways, ranging from protein-level enrichments for tagged Ub prior to digestion and analysis^15,16^, down to enrichment of characteristic peptidic ubiquitin remnants that are left behind following protein digestion^12,17,18^. Perhaps the most common method for ubiquitination profiling by proteomics is through the use of a GG-K remnant antibody^18^, where samples are digested using trypsin, which cleaves amide bonds that are C-terminal to basic residues such as arginine and lysine. When a ubiquitinated protein is digested in this manner, it leaves a characteristic diglycyl remnant on the lysine residues that previously had been sites for ubiquitination. This diglycyl remnant is then the target for GG-K remnant antibodies that are often bound to resins or magnetic beads, enabling facile enrichment of GG-K peptides (Figure 1). While convenient and robust, these antibodies are known to have sequence biases for flanking residues of the enriched peptides^18,19^ limiting their overall utility. Further, these antibodies detect only lysine sidechain modifications, thereby missing noncanonical ubiquitination sites at protein *N*- termini and on other amino acid sidechains. To circumvent these difficulties, several groups have developed novel strategies for ubiquitin enrichment and downstream proteomics. One such strategy, UbiSite, was developed by Blagoev and colleagues to enable enrichment of non-lysine ubiquitination and to discern Ub scars from other Ub-like proteins (such as NEDD8 and ISG15)^11^. This approach uses a different antibody to specifically recognize ubiquitylation sites via the 13-residue scar from the C-terminus of ubiquitin left on a protein after cleavage with LysC.

**Figure 1.**
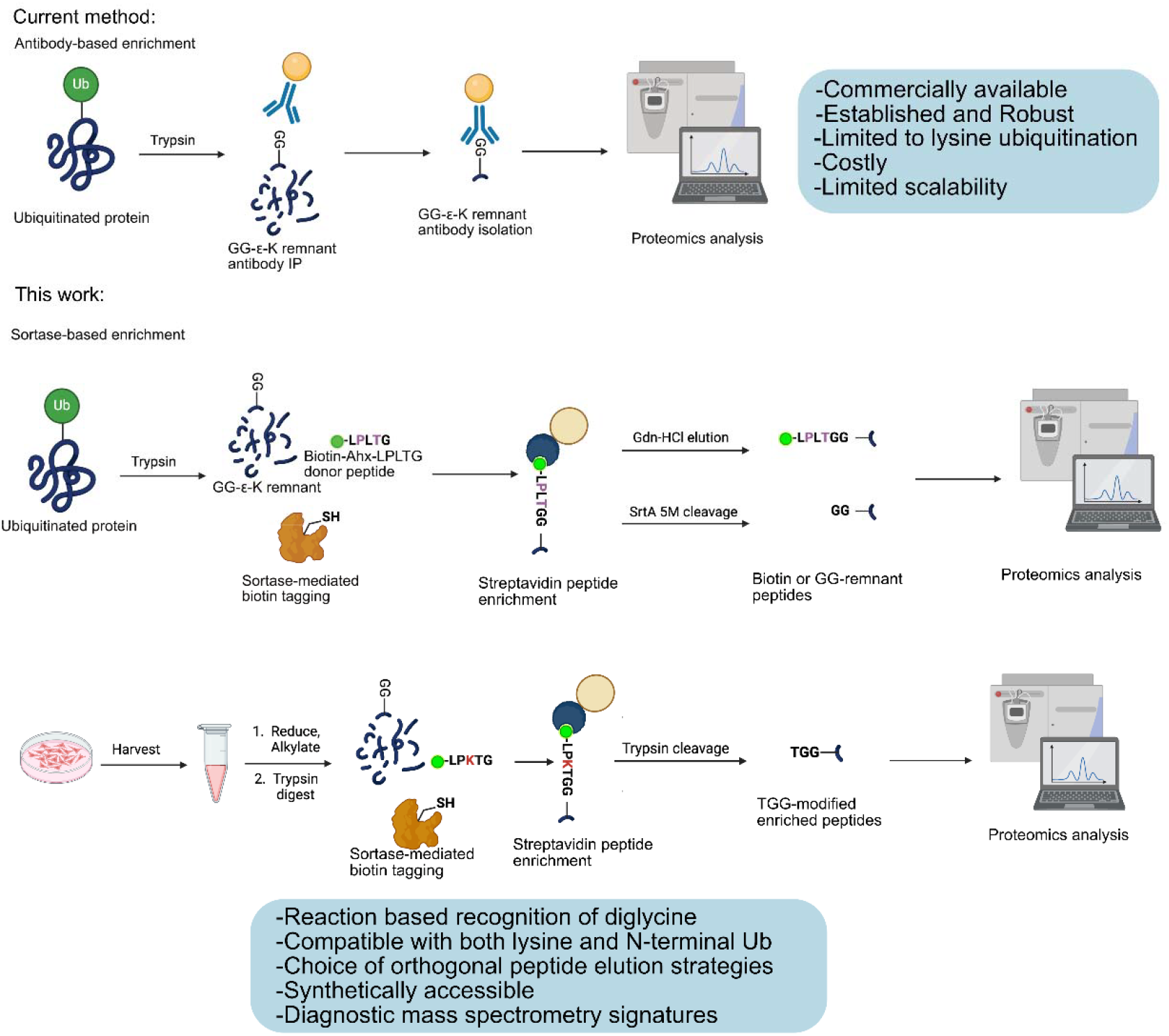
Methods for ubiquitin enrichment and proteomics. Top: General workflow for antibody-based enrichment strategies of peptides with ubiquitin diglycyl lysine (GG-K) remnant. Bottom: New workflows for sortase-based enrichment strategies presented in this work.

In addition to antibody-driven approaches, other workflows for the enrichment of ubiquitinated proteins for proteomics studies have been reported. Two of these methods specifically involve the blocking of unmodified lysine residues and peptide *N*-termini, followed by the removal of Ub groups using nonspecific DUBs, with Sun *et. al*. drawing specifically on click chemistry to enrich for peptides after Ub removal^20^. Alternatively, Stes *et. al*. leveraged the reactivity of the free lysines to conjugate a Boc protecting group and observed the changes in chromatographic properties of the peptides^21^. While these creative methods bypass the need for sequence biased and context-specific antibodies, they rely on extensive chemical modifications of the samples and complex analysis of the modified peptides. To circumvent some of these pitfalls, our lab turned to biochemical methods to enrich for ubiquitinated peptides for downstream proteomic analysis.

Based on our previous work^22^, we reasoned that the diglycine scar generated during proteolysis of ubiquitinated proteins could serve not only as an analytical signature of ubiquitination, but also as a chemically useful intermediate. Sortase A is a bacterial transpeptidase enzyme that has been employed in countless bioconjugation efforts and is dependent on an *N*-terminal glycine residue (but is commonly used with several consecutive glycine residues) as an “acceptor” for the “donor” protein bearing a sortase-compatible LPXTG motif on the tagged protein for conjugation. More recently Sortase A is gaining traction in its ability to be utilized for site-specific conjugation of ubiquitin and ubiquitin-like modifiers^22–24^, suggesting that ubiquitin-derived diglycine motifs could provide intrinsic acceptors for post-digest enzymatic labeling. Such a strategy would convert information encoded by ubiquitination into a new covalent handle without requiring genetic manipulation of ubiquitin or recognition by a remnant-specific antibody.

Here, we establish this concept using engineered Sortase A to remodel ubiquitin-derived diglycine remnants with functionalized peptide donors. We first define the reaction on synthetic and recombinant ubiquitinated substrates, then extend it to peptides generated from mammalian cell lysates. The resulting conjugates generate characteristic fragmentation signatures that aid their mass-spectrometric identification. We further exploit the modularity of the sortase donor to encode orthogonal release mechanisms: reversible sortase chemistry restores the diglycine-modified peptide, whereas incorporation of an internal trypsin cleavage site converts enriched peptides into a compact TGG-modified product. Together, these studies demonstrate that a proteolytic PTM scar can be repurposed as a substrate for chemoenzymatic manipulation and affinity enrichment.

## Results and Discussion

### Ubiquitin-Derived Diglycine Scars Are Substrates for Sortase-Mediated Remodeling

Sortase-mediated conjugation is typically performed on substrates containing 3+ glycine residues at the *N*-terminus followed by the target of interest. While this has been shown to be the most effective substrate for sortylation, we have previously demonstrated sortase-mediated conjugation can take place on a single *N*-terminal glycine^22^, and the Lang group has demonstrated robust sortase-mediated ubiquitination using their non-canonical amino acid diglycyl lysine (GG-K) as the glycyl acceptor^23^. Given these data, we hypothesized that GG-K, both alone and within a longer peptide, would be suitable substrates for sortase-mediated conjugation. To test this, we initially turned to a peptide-based system to investigate the conjugation of a biotin-labeled peptide bearing the sortase recognition motif (Biotin-Ahx-LPLTG) to a synthesized GGK peptide *in vitro* (Figure 2). To synthesize the GGK acceptor peptide, we performed a simple amide coupling between (Boc GG and Boc-Lys-tBu) before a purification followed by an acidic global deprotection of the peptide (Supplemental Scheme 1, Supplemental Fig. 1). For the Biotin-Ahx-LPLTG peptide (Figure 1), we utilized standard solid-phase peptide synthetic methods. Upon deprotection and purification, we observed the anticipated masses for each of the peptides by LC-MS (Supplemental Fig. 2). We initially performed a sortase assay *in vitro* using the SrtA 7M mutant^26^ and observed successful bioconjugation between the two peptides, yielding the Biotin-Ahx-LPLTGGK product peptide that could be detected by LC-MS (Fig. 2, Supplemental Fig. 3). To better understand the kinetics of this conjugation and for optimization in later experiments, time points were taken at 0 hours, 4 hours, and 16 hours, and with the 16-hour time point used in subsequent workflows based on the highest conjugation of the two peptides (Figure 2).

**Figure 2.**
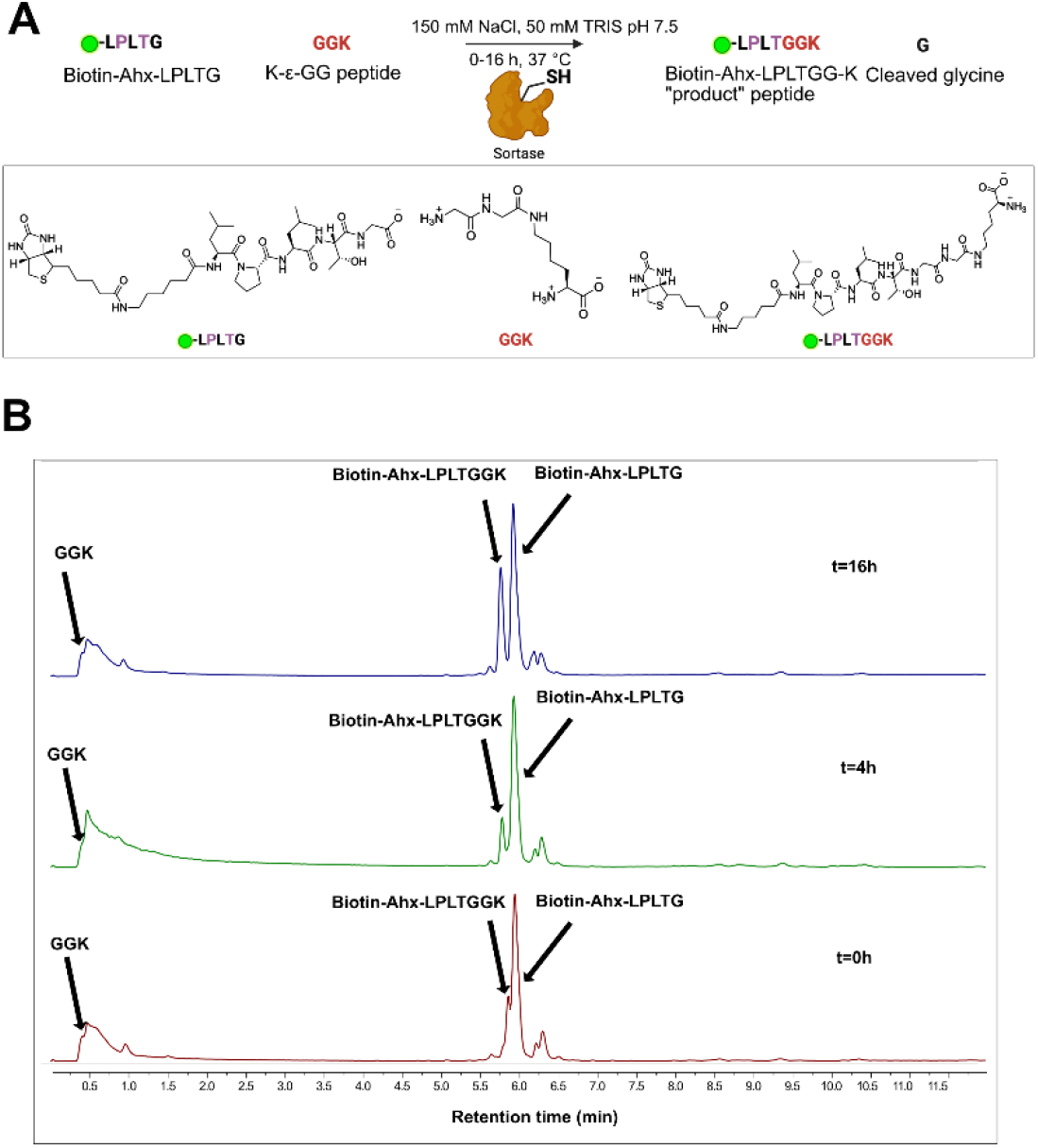
Chemoenzymatic remodeling of a ubiquitin-remnant mimic. A) General approach for sortase conjugation of Biotin-Ahx-LPLTG to GGK acceptor peptide (top). Chemical structures and shorthand schematics for peptide substrates and products (bottom). B) Time course experiment of

### Sortase Remodels Diglycine Scars on Defined Ubiquitinated Proteins

Following confirmation that the peptide-based conjugation assay was successful, we tested the bioconjugation ability of sortase on peptides generated from ubiquitinated protein samples. We employed purified K63 and K48-linked diubiquitin, which were readily prepared and purified *in vitro* using standard protocols^22,27^. Following purification of the di-ubiquitins, the proteins were digested using either trypsin or a trypsin/LysC combination protease for a minimum of 6 hours before analysis of the peptides by LC-QTOF (Fig. 3A, schematic). We observed the diglycyl remnant from the distal ubiquitin and obtained good coverage of the peptides bearing the diglycyl mark at the correct position based on the diubiquitin starting material (Figure 3A, Supplemental Fig. 4), demonstrating that the protease digest and data analysis methods were compatible and sensitive enough to detect the diglycyl mark.

**Figure 3.**
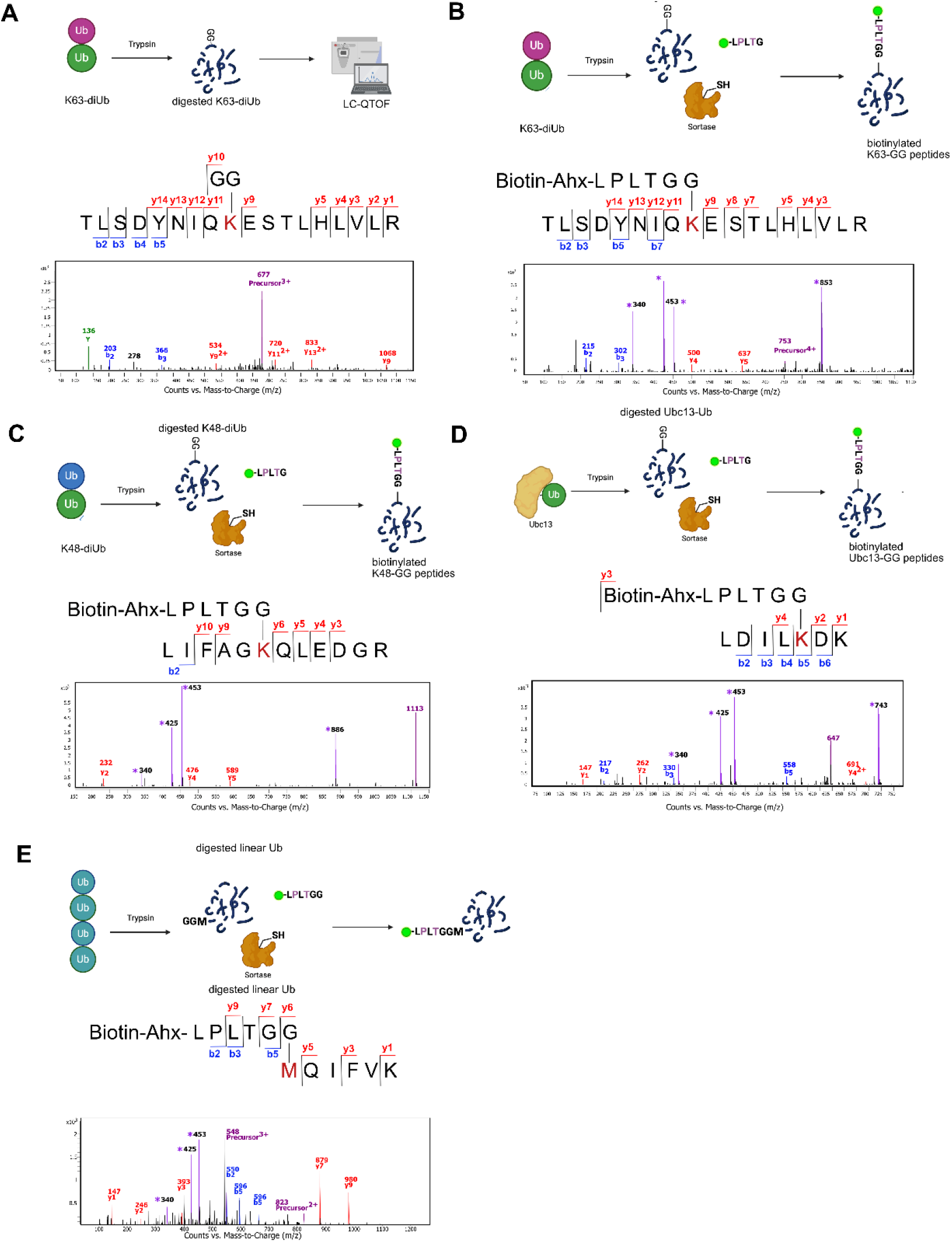
Sortase-mediated remodeling of ubiquitin-derived diglycine remnants. A) Schematic showing general workflow for digestion of K63-linked diubiquitin and mass spec detection. Associated b and y ions showing diglycyl remnant at position K63 are indicated along the linear peptide sequence with a representative spectrum shown. B) Schematic showing general workflow for digestion and sortase-mediated biotinylation of diglycyl remnant peptides at position K63 on ubiquitin. Linear peptide sequence is labelled with identified b and y ions showing Biotinylated adduct. C) Schematic, linear sequences, and representative mass spectra of K48-diubiquitin digested peptides after biotinylation. Purple asterisks indicate Biotin-Ahx-LPLTGG and branched peptide fragments not recognized in the initial data processing. D) Schematic, linear sequences, and representative mass spectra of digested Ubc13-Ub showing biotinylated peptides. Purple asterisks indicate Biotin-Ahx-LPLTG and branched peptide fragments not recognized in the initial data processing. E) Schematic, peptide sequence, and representative mass spectrum of biotinylated peptide from linear polyubiquitin showing *N*-terminal ubiquitination.

Given these data in combination with the workflow established for peptide biotinylation, we next tested if we could observe biotin peptide conjugation to digly peptides from digested ubiquitinated proteins via LC-QTOF. To do this, we performed a tryptic digest of the diubiquitin, followed by sortase-mediated ligation of the biotin peptide onto the digested diubiquitin peptides, then analyzed the products by LC-QTOF (Fig. 3B, 3C). To prevent any undigested sortase protein from being subjected to the peptide column on the LC-QTOF, samples were initially subjected to 10% formic acid precipitation, and the soluble portion was analyzed, but more complete coverage of the peptides was observed when the reaction mixture was subjected to a second tryptic digest prior to analysis. By LC-QTOF, we were pleased to see peptides corresponding to the biotinylated versions of peptides that previously bore diglycyl remnants, with conversions up to 82%. In parallel, we also tested this strategy on an *in vitro* ubiquitinated E2 ubiquitin conjugating enzyme Ubc13. In testing this strategy, we observed autoubiquitination at Ubc13’s own lysine 82 and lysine 92 (Supplemental Fig. 5). Crucially, the ubiquitination of K92 has been noted in many high-throughput datasets, and the UbcH8-mediated ISGylation of this position has been reported to inhibit Ubc13’s ability to bind ubiquitin^28,29^. Upon testing our sortase-mediated biotinylation strategy on digested Ubc13-Ub, we observed successful biotinylation of the diglycyl remnant on K92, as demonstrated in Fig. 3D.

To test the applicability of this technique for noncanonical ubiquitination sites, we next tested whether we could conjugate our biotin peptide to the *N-*terminus of ubiquitinated proteins. For this scenario, we utilized linear tetraubiquitin, which can be expressed *in vitro* as a fusion protein, and purified according to Chang et. al.^30^. Upon trypsin digest and sortase-mediated biotinylation, we successfully detected peptides consistent with biotinylation of the *N*-terminal diglycyl remnant, (Fig. 2E) demonstrating that this strategy is applicable for non-lysine ubiquitination sites, unlike antibody based enrichment.

### Chemoenzymatic Remodeling Generates Diagnostic Fragmentation Signatures in Complex Proteomes

Following these initial proof of concept studies using recombinant proteins, we next tested whether we could detect biotinylated peptides from endogenous ubiquitination sites in more complex samples. To do this, we adapted a standard proteomics workflow to our new strategy using cell lysates from HEK293T cells (Fig. 4A). Cells were harvested and proteins subjected to reduction, alkylation, and trypsin digestion prior to sortase-mediated biotinylation (Fig. 4A). When detecting the biotinylated peptides, it is important to note that the product of biotinylation on diglycyl-lysine remnants generates a branched peptide that is subject to additional fragmentation during ionization, resulting in diagnostic ions (Fig. 4B). While this branched peptide may elude standard proteomics workflows, the diagnostic ions (Fig. 4B) can be used to narrow the search query in the datasets. Using these diagnostic masses in combination with lysine adducts generated by sortase, we detected several known and high confidence lysine ubiquitination sites (Table 1), with strong spectral data (Fig. 4C/D). Additionally, by generating a unique search database of *N*- terminally labelled peptides, we were able to identify sites of *N*-terminal ubiquitination in unenriched mammalian cell lysates (Table 1, Fig. 4E). Taken together, these data show the outlined sortase-mediated biotinylation strategy enables detection of ubiquitin sites in mammalian cell lysates.

**Figure 4.**
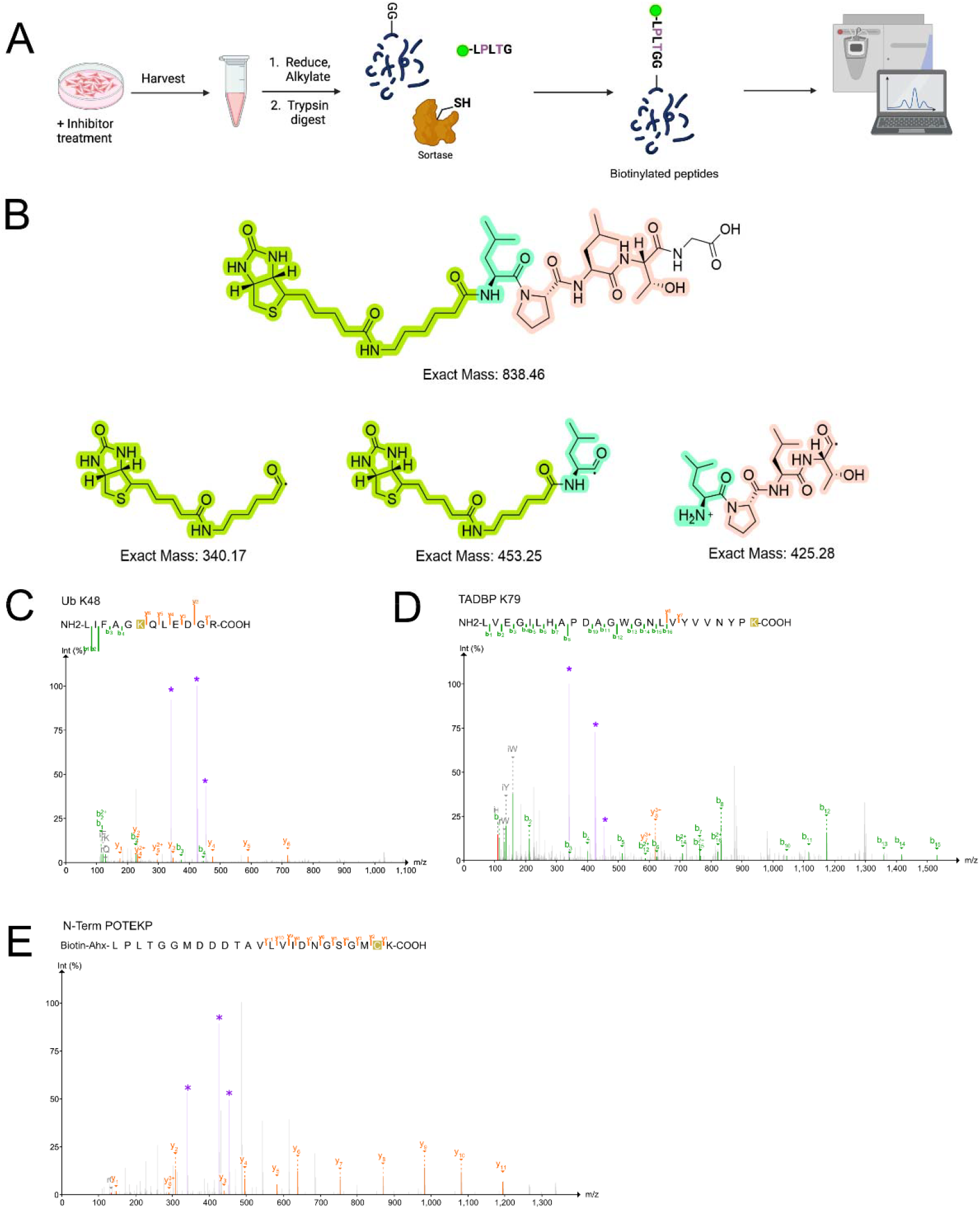
Diagnostic fragmentation of remodeled ubiquitin-remnant peptides in mammalian cell lysates. A) Workflow for biotinylation of ubiquitinated proteins in mammalian cell lysates. B) Diagnostic ions from branched peptide Biotin-Ahx-LPLT fragmentation. Representative spectra from detected ubiquitination sites for Ubiquitin K48 C) and TADBP K79 (D) and POTEKP *N*-term (E). Diagnostic ions are marked with *.

**Table 1.**
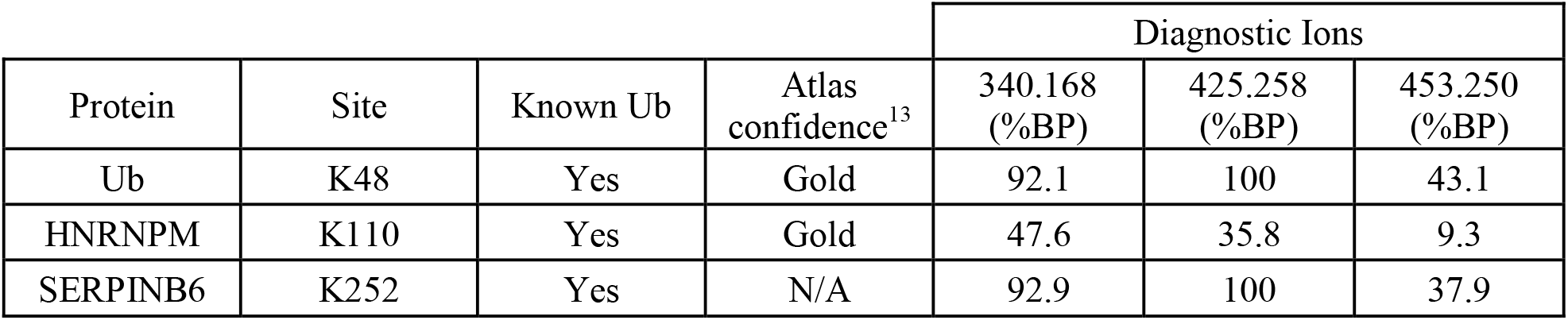
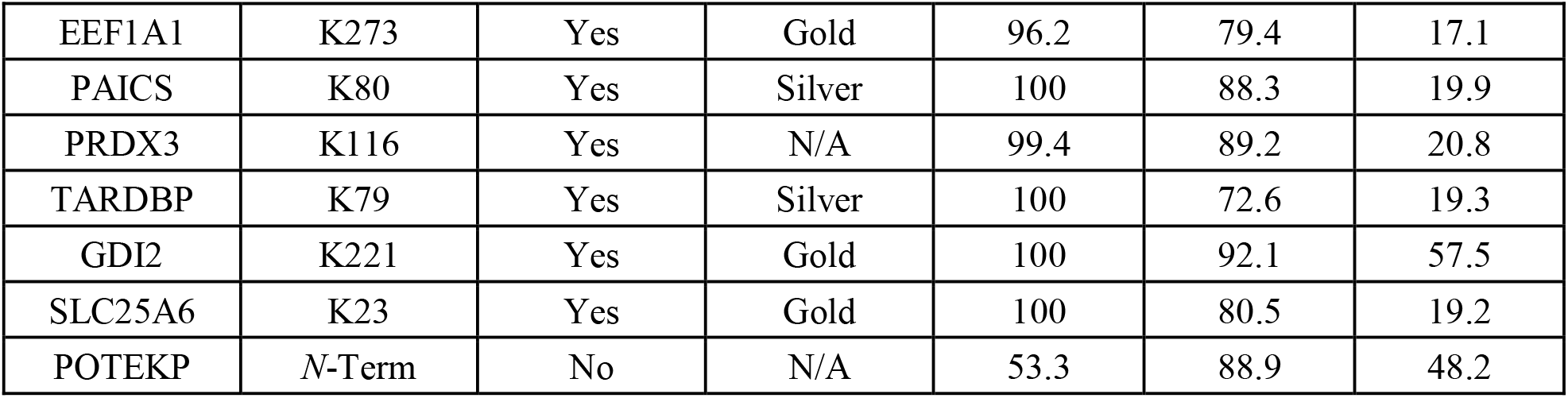
Identified Ubiquitin Sites and Diagnostic Ions.

Following successful chemoenzymatic tagging of the peptides, we next tested whether the biotinylated *iso*peptide would serve as an effective handle for affinity enrichment. Leveraging the tight binding affinity between biotin and streptavidin, we selected a streptavidin bound to agarose beads for our enrichment strategy. Further, we envisioned that we could leverage two potential elution strategies to release the peptides from the resin (Fig. 5A). Using a classic elution with guanidinium hydrochloride (GdnHCl) at low pH, we hypothesized that this method would recover the biotinylated peptides from the initial workflow. Initially testing this strategy on *in vitro* purified, digested, and biotinylated K48-linked diubiquitin, we successfully enriched the peptide containing K48 from the mixture and saw the subsequent release of the biotinylated peptide upon elution using GdnHCl (Fig. 5B). Complementing this workflow, we also tested an elution strategy that leveraged the reversibility of a different mutant of sortase^22^, SrtA 5M^31^, to hydrolyze the branched biotinylated peptides, releasing the diglycyl-containing peptides. Incubation of the biotinylated peptides with SrtA 5M indeed released the diglycyl modified peptides (Fig. 5C), and we observed better coverage across peptides following elution with sortase. In consideration of more standard proteomics workflows that utilize trypsin, we also developed a bifunctional biotin peptide, Biotin-Ahx-LPKTG (Fig. 5D). This bifunctional peptide retained the “LPXTG” motif rendering it sortase compatible with the additional benefit of containing a trypsin-labile internal lysine residue, enabling a sortase-independent strategy for release of the enriched peptides from the streptavidin beads. Eluting via trypsin cleavage, we observe “TGG” modified peptides due to the trypsin cleavage site appearing N-terminal to the threonine residue in Biotin-Ahx-LPKTG (Fig. 5E), providing a unique handle for tracking the +215 Da remnant. By adapting this strategy to the environment of mammalian cell lysates (Fig. 5F), we successfully enriched several known endogenous ubiquitination sites (Fig. 5G/H), demonstrating enrichment of ubiquitin derived peptides, in a sortase-mediated manner, from complex biological samples.

**Figure 5.**
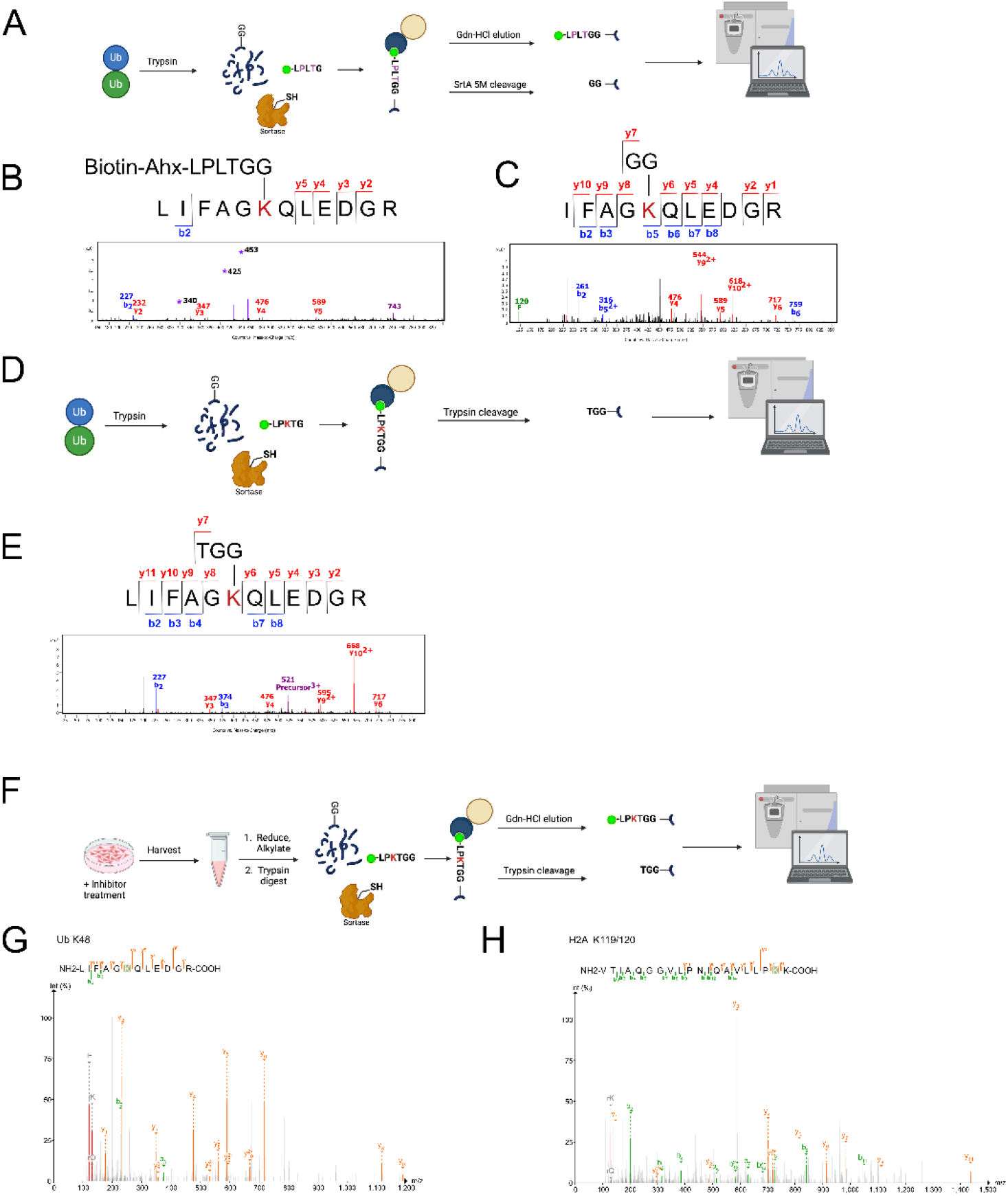
Programmable affinity enrichment and release of remodeled ubiquitin-remnant peptides. A) Schematic showing general workflow for ubiquitinated protein digest, sortase-mediated biotinylation using Biotin-Ahx-LPLTG peptide, enrichment, and elution from beads. Peptides can be eluted via incubation with 6M GdnHCl at pH 1.5 to release biotinylated diglycyl lysine sites (top) or cleaved from the resin leveraging SrtA 5M (bottom). B) Representative MS2 spectrum showing GdnHCl elution of biotinylated K48 ubiquitin peptide. Diagnostic masses of 340, 425, and 453 Da are indicated with purple asterisks. C) Representative MS2 spectrum showing SrtA 5M-mediated elution of biotinylated K48 ubiquitin peptide. D) Schematic showing general workflow for ubiquitinated protein digest, sortase-mediated biotinylation using Biotin-Ahx-LPKTG peptide, enrichment, and trypsin-mediated release of enriched peptides from beads. E) Representative MS2 spectrum showing trypsin elution of TGG-modified K48 ubiquitin peptide following enrichment. F) Workflow for enrichment of ubiquitinated peptides from mammalian cell lysates using bifunctional Biotin-Ahx-LPKTG peptide. Representative spectra of identified TGG remnant peptides from Ubiquitin (G) and H2A (H)

The LPKTG donor demonstrates that the chemistry of the enrichment reagent can be used to program the molecular form in which captured ubiquitin sites are returned for analysis. Whereas the LPLTG reagent retains the complete sortase-derived branch unless hydrolyzed enzymatically, introduction of a single lysine into the donor creates a protease-cleavable junction that converts the affinity-tagged conjugate into a compact TGG remnant. Thus, recognition, enrichment, release, and MS encoding can be independently engineered through donor-peptide design.

## Discussion

This work establishes proteolytic ubiquitin remnants as substrates for post-digest chemoenzymatic remodeling. Rather than treating the diglycine remnant solely as a diagnostic mass shift, we exploit its intrinsic chemo-enzymatic reactivity as a sortase acceptor to install functionality directly at sites derived from ubiquitination. In this way, proteolysis converts a post-translational modification into a chemically tractable intermediate that can be manipulated before mass-spectrometric analysis.

The approach operates across increasing levels of molecular complexity, from synthetic diglycine-containing peptides to defined ubiquitinated proteins and peptides generated from mammalian cell lysates. Importantly, the strategy is not restricted to lysine-linked diglycine remnants: *N*-terminal diglycine generated from linear ubiquitin is also accepted by sortase. The ligation products additionally exhibit characteristic fragmentation behavior, providing an independent analytical signature of the installed chemistry. Finally, modification of the LPXTG donor sequence enables control over how enriched peptides are released and subsequently represented in the mass spectrum.

This chemistry is orthogonal to conventional remnant immunoaffinity enrichment. Whereas antibody-based workflows rely on molecular recognition of a diglycyl-lysine epitope, the present approach uses the diglycine motif as a reaction substrate. Consequently, capture and release functionality can be encoded into the sortase donor rather than being intrinsic to the recognition reagent. The extent to which local peptide sequence influences sortase reactivity in proteome-scale applications remains to be established and will require systematic comparison across larger datasets but it’s important to note that the widely-used GG-K exhibit a flanking sequence bias toward leucines and acidic residues^18,19^.

Recent work from Komander and colleagues independently demonstrated that diglycine-containing products generated from non-protein ubiquitination can be derivatized using sortase for metabolomic analysis^25^. The distinct substrate classes and analytical objectives of that study and the present work highlight a broader chemical feature of ubiquitin-derived diglycine motifs: following appropriate proteolytic processing, they provide enzymatically addressable handles that can be remodeled for downstream analysis. Here, we exploit this chemistry directly on protein-derived ubiquitination sites and extend it to affinity enrichment, alternative release strategies, and proteomic detection. When considered alongside the strategy presented here, the complementarity of Sortase to ubiquitin systems proves a powerful tool to continue to unravel the mysteries of ubiquitination. We hope that others in the field are equally inspired by these approaches and continue to develop new strategies to combat the challenges in ubiquitin detection, further expanding the scope of what is currently understood about the modification.

More broadly, these studies illustrate how a proteolytic PTM scar can be treated as a synthetic intermediate rather than solely as an analytical signature. By exploiting the intrinsic reactivity of ubiquitin-derived diglycine motifs, sortase-mediated remodeling couples molecular recognition to installation of user-defined chemical functionality. The modularity of the donor peptide further enables affinity capture, enzymatic release, and encoding of distinctive mass-spectrometric signatures within a common reaction framework. We anticipate that this scar-directed chemoenzymatic strategy will provide a foundation for new approaches to manipulate and analyze ubiquitination and may motivate analogous strategies in which proteolysis-generated signatures of other biomolecular modifications are repurposed for downstream chemistry.

## Acknowledgments

We thank members of the Burslem lab for comments and intellectual discussion over the course of this study. We would also like to acknowledge Dr. Altaf Najar for his assistance with the mass spectrometry data acquisition. This work was supported by the National Institute of General Medical Sciences (R35GM142505) to G.M.B., the NIH Chemistry Biology Interface Training Grant (T32 GM133398) to N.R.R. and K.B., and an NSF Graduate Research Fellowship to M.N.

## Notes

### Competing Interest Statement

The authors have declared no competing interest.

### Summary of Updates

New data added - reformatted and text editied

