## Supplementary Information for "Ubiquitin-Derived Diglycine Remnants as Chemical Handles for Chemoenzymatic Proteomics"

### Supplemental Materials and Methods

#### General Methods

Proteins generated *in vitro* were digested into peptides using Trypsin. Proteins were dissolved or dialyzed into 100 mM tri-ethyl ammonium bicarbonate (TEABC) buffer, pH 7.5, and protease was added to the sample in a 1:20 protease:substrate ratio. Samples were digested at 37 °C for 6-16 hours. Following digestion, protease was quenched by adjusting samples to pH 3.0 using formic acid, and excess reaction buffer was removed under reduced pressure. Samples were resuspended in 50 mM TEABC and any precipitated proteins removed by centrifugation for 10 minutes at 21,300 rcf.

#### Generation of constructs

pet30b-7M SrtA and pet30b-5M SrtA plasmids were a generous gift from Hidde Ploegh (Addgene plasmids # 51141 and # 51140). pET3a-hUb plasmid was a generous gift from Chris Hill (Addgene plasmid #61937). pDEST17-Cdc34a was a generous gift from Wade Harper (Addgene plasmid # 18674). pETSUMO-hUbc13 was a generous gift from Cynthia Wolberger (Addgene plasmid # 51131). pET28aLIC-UEV1 was a generous gift from Cheryl Arrowsmith (Addgene plasmid # 25619). pET28-UBE1 was a generous gift from Deschaies lab. GST-Ub4 was a gift from James Hurley (Addgene plasmid # 171421).

#### Recombinant Protein Sequences

UBE1
MGSSHHHHHHSSGLVPRGSHMSSSPLSKKRRVSGPDPKPGSNCSPAQSALSEVSSVPTNGMAKN
GSEADIDESLYSRQLYVLGHEAMKMLQTSSVLVSGLRGLGVEIAKNIILGGVKAVTLHDQGTTQ
WADLSSQFYLREEDIGKNRAEVSQPRLAELNSYVPVTAYTGPLVEDFLSSFQVVVLTNSPLEAQL
RVGEFCHSRGIKLVVADTRGLFGQLFCDFGEEMVLTDSNGEQPLSAMVSMVTKDNPGVVTCLD
EARHGFETGDFVSFSEVQGMIQLNGCQPMEIKVLGPYTFSICDTSNFSDYIRGGIVSQVKVPKKISF
KSLPASLVEPDFVMTDFAKYSRPAQLHIGFQALHQFCALHNQPPRPRNEEDATELVGLAQAVNA
RSPPSVKQNSLDEDLIRKLAYVAAGDLAPINAFIGGLAAQEVMKACSGKFMPIMQWLYFDALEC
LPEDKEALTEEKCLPRQNRYDGQVAVFGSDFQEKLSKQKYFLVGAGAIGCELLKNFAMIGLGCG
EGGEVVVTDMDTIEKSNLNRQFLFRPWDVTKLKSDTAAAAVRQMNPYIQVTSHQNRVGPDTER
IYDDDFFQNLDGVANALDNIDARMYMDRRCVYYRKPLLESGTLGTKGNVQVVIPFLTESYSSSQ
DPPEKSIPICTLKNFPNAIEHTLQWARDEFEGLFKQPAENVNQYLTDSKFVERTLRLAGTQPLEVL
EAVQRSLVLQRPQTWGDCVTWACHHWHTQYCNNIRQLLHNFPPDQLTSSGAPFWSGPKRCPHP
LTFDVNNTLHLDYVMAAANLFAQTYGLTGSQDRAAVASLLQSVQVPEFTPKSGVKIHVSDQEL
QSANASVDDSRLEELKATLPSPDKLPGFKMYPIDFEKDDDSNFHMDFIVAASNLRAENYDISPAD
RHKSKLIAGKIIPAIATTTAAVVGLVCLELYKVVQGHQQLDSYKNGFLNLALPFFGFSEPLAAPR
HQYYNQEWTLWDRFEVQGLQPNGEEMTLKQFLDYFKTEHKLEITMLSQGVSMLYSFFMPAAKL
KERLDQPMTEIVSRVSKRKLGRHVRALVLELCCNDESGEDVEVPYVRYTIR

Cdc34
MSYYHHHHHHLESTSLYKKAGSAAAPFTMARPLVPSSQKALLLELKGLQEEPVEGFRVTLVDEG
DLYNWEVAIFGPPNTYYEGGYFKARLKFPIDYPYSPPAFRFLTKMWHPNIYETGDVCISILHPPVD

DPQSGELPSERWNPTQNVRTILLSVISLLNEPNTFSPANVDASVMYRKWKESKGKDREYTDIIRK
QVLGTKVDAERDGVKVPTTLAEYCVKTKAPAPDEGSDLFYDDYYEDGEVEEEADSCFGDDEDD
SGTEES

His-SUMO-hUbc13
MGHHHHHHPSGVKTENNDHINLKVAGQDGSVVQFKIKRHTPLSKLMKAYCERQGLSMRQIRFR
FDGQPINETDTPAQLEMEDEDTIDVFQQQTGGMAGLPRRIIKETQRLLAEPVPGIKAEPDESNARY
FHVVIAGPQDSPFEGGTFKLELFLPEEYPMAAPKVRFMTKIYHPNVDKLGRICLDILKDKWSPAL
QIRTVLLSIQALLSAPNPDDPLANDVAEQWKTNEAQAIETARAWTRLYAMNNI

UEV1
MGSSHHHHHHSSGLVPRGSTGVKVPRNFRLLEELEEGQKGVGDGTVSWGLEDDEDMTLTRWT
GMIIGPPRTIYENRIYSLKIECGPKYPEAPPFVRFVTKINMNGVNSSNGVVDPRAISVLAKWQNSY
SIKVVLQELRRLMMSKENMKLPQPPEGQCYSN

Ub WT
MQIFVKTLTGKTITLEVEPSDTIENVKAKIQDKEGIPPDQQRLIFAGKQLEDGRTLSDYNIQKESTL
HLVLRLRGG

UbK0
MQIFVRTLTGRTITLEVEPSDTIENVRARIQDREGIPPDQQRLIFAGRQLEDGRTLSDYNIQRESTL
HLVLRLRGG

SrtA 5M
MQAKPQIPKDKSKVAGYIEIPDADIKEPVYPGPATREQLNRGVSFAEENESLDDQNISIAGHTFID
RPNYQFTNLKAAKKGSMVYFKVGNETRKYKMTSIRNVKPTAVEVLDEQKGKDKQLTLITCDDY
NEETGVWETRKIFVATEVKLEHHHHHH

SrtA 7M
MQAKPQIPKDKSKVAGYIEIPDADIKEPVYPGPATREQLNRGVSFAKENQSLDDQNISIAGHTFID
RPNYQFTNLKAAKKGSMVYFKVGNETRKYKMTSIRNVKPTAVEVLDEQKGKDKQLTLITCDDY
NEETGVWETRKIFVATEVKLEHHHHHH

GST-Ub4
MSPILGYWKIKGLVQPTRLLLEYLEEKYEEHLYERDEGDKWRNKKFELGLEFPNLPYYIDGDVK
LTQSMAIIRYIADKHNMLGGCPKERAEISMLEGAVLDIRYGVSRIAYSKDFETLKVDFLSKLPEM
LKMFEDRLCHKTYLNGDHVTHPDFMLYDALDVVLYMDPMCLDAFPKLVCFKKRIEAIPQIDKY
LKSSKYIAWPLQGWQATFGGGDHPPKSDLVPRGSPGIHMQIFVKTLTGKTITLEVEPSDTIENVK
AKIQDKEGIPPDQQRLIFAGKQLEDGRTLSDYNIQKESTLHLVLRLRGGMQIFVKTLTGKTITLEV
EPSDTIENVKAKIQDKEGIPPDQQRLIFAGKQLEDGRTLSDYNIQKESTLHLVLRLRGGMQIFVKT
LTGKTITLEVEPSDTIENVKAKIQDKEGIPPDQQRLIFAGKQLEDGRTLSDYNIQKESTLHLVLRLR
GGMQIFVKTLTGKTITLEVEPSDTIENVKAKIQDKEGIPPDQQRLIFAGKQLEDGRTLSDYNIQKES
TLHLVLRLRGG

**Protein expression and purification**

Ubiquitin was expressed and purified as previously described[1]. Briefly, pET3a-hUb was expressed in Rosetta2 *E. coli* cells in Lura-Bertani (LB) media supplemented with Kanamycin. Cells were grown while shaking in an incubator at 37 °C until they reached an $OD_{600}$=0.6-0.8. Cells were induced with 0.5 mM isopropyl β-D-1-thiogalactopyranoside (IPTG) and expressed at 37 °C for 3-5 hours before harvesting by centrifugation. Cells were resuspended in a minimal amount of lysis buffer (10 mM TRIS pH 8.0, 150 mM NaCl, protease inhibitor cocktail pellet [Thermo]), and pellets were stored at -80 °C until purification. Cell

pellets were thawed before lysis by sonication on ice, and lysates were cleared by centrifugation for 45 minutes at 13,000 x g. The soluble fraction was filtered through a 30 kDa MWCO centrifugation filter (Pierce or Amicon) before concentration in a 3 kDa MWCO centrifugal filter. The concentrated material was centrifuged for 10 min x 21300 rcf before purification by size-exclusion chromatography (SEC) on a Superdex HiLoad 16/600 S75 column (Cytiva), and purified protein was stored at -80 °C until future use.
UBE1 was expressed and purified as previously described[2]. Briefly, Rosetta 2 *E. coli* cells were grown in LB media supplemented with Kanamycin. Cells were grown at 37 °C while shaking until $OD_{600}$ = 0.6. The shaker temperature was adjusted to 16 °C and cells were induced with 0.5 mM IPTG and expressed overnight before collecting cells via centrifugation. Cell pellets were resuspended in 50 mM TRIS pH 8.0, 150 mM NaCl, 1 mM DTT, and 1 protease inhibitor cocktail pellet before storing at -80 °C. Cells were sonicated and lysates cleared as described above. The soluble fraction was incubated with NiNTA-Agarose resin (Qiagen) at 4 °C for 1 hour before purification. Resin was washed with resuspension buffer and eluted with resuspension buffer supplemented with 100 mM imidazole. Protein was dialyzed against 50 mM TRIS pH 8.0, 150 mM NaCl, and 1 mM DTT overnight at 4 °C before purification by SEC using a Superdex HiLoad 16/600 S200 column (Cytiva). Final collected protein was flash frozen in liquid nitrogen and stored at -80 °C until use.
UEV1 and Ubc13 were expressed and purified as described[3]. Briefly, Rosetta 2 *E. coli* cells were grown in LB media supplemented with Kanamycin (UEV1) or Ampicillin (Ubc13) and induced at $OD_{600}$ = 0.6 with 0.5 mM IPTG. Cells expressed E2s at 16 °C overnight and were collected by centrifugation. Cell pellets were resuspended in 50 mM TRIS pH 8.0, 300 mM NaCl, and 1 mM DTT supplemented with a protease inhibitor pellet.

Linear tetraubiquitin was expressed and purified as described[4]. Briefly, Rosetta 2 *E. coli* cells were grown in LB media supplemented with Ampicillin and were induced at $OD_{600}$ = 0.6 with 0.2 mM IPTG. Cells expressed at 16 ºC overnight and were harvested by centrifugation and stored at -80 ºC until needed. For purification, pellet was resuspended in 50 mM HEPES (7.5), 300 mM NaCl, 1 mM DTT, and 1 protease inhibitor cocktail. Cells were lysed by sonication and lysates cleared as described above. The soluble fraction was incubated with Glutathione Sepharose 4B resin at 4 ºC for 2 hours, followed by 30 minutes at room temperature. Resin was washed with lysis buffer and linear tetraubiquitin eluted with lysis buffer containing 50 mM glutathione.

**Generation and purification of diubiquitin constructs**
Diubiquitin constructs were generated and purified as previously described[5]. Briefly, polyubiquitin synthesis reactions were conducted using 1 uM UBE1, 5 uM E2 (employing His-SUMO-Ubc13 and UEV1 for K63-linked diubiquitin and Cdc34 for K48-linked diubiquitin), 400 uM Ubiquitin, 50 mM Tris pH 8.0, 4 mM MgCl2, and 2 mM ATP. Reactions were conducted at 37 °C in a benchtop shaker for 1.5 hours (for K63-linked diubiquitin) or 4 hours (for K48-linked diubiquitin) before flash freezing and storage at -80 °C. For purification, reactions were thawed and concentrated using Pierce or Amicon ultracentrifugation filters (3 kDa MWCO) to 100 uL. Reaction was diluted 10-fold into polyubiquitin ion exchange buffer A (50 mM sodium acetate pH 4.5) and precipitate cleared by centrifugation at 21100 x rcf for 10 minutes. Soluble polyubiquitin was purified on a Capto S 1 mL cation exchange column on a gradient of 0-50% buffer B (50 mM sodium acetate pH 4.5, 1 M NaCl) over 340 mL before a final wash of 100% buffer B (5 mL). Fractions were analyzed by SDS-PAGE and purified diUb fractions were pooled, concentrated using Pierce ultracentrifugation filters, and flash frozen using liquid nitrogen before storage at -80 °C.

**Generation and purification of mono-ubiquitinated SUMO-Ubc13**
Mono-ubiquitinated SUMO-Ubc13 (denoted as Ubc13-Ub) was generated *in vitro* in reactions containing 1 uM UBE1, 100 uM SUMO-Ubc13, 12 uM UbK0, 4 mM MgCl2, 50 mM TRIS (pH 8.0), and 2 mM ATP. Reactions were conducted at 37 °C on a benchtop shaker overnight and observed by SDS-PAGE. Reactions were centrifuged at 21100 rcf x 10 minutes and purified by SEC on an S75 gel filtration column pre-equilibrated in 300 mM NaCl, 50 mM TRIS pH 8.0, and 1 mM DTT. Fractions containing Ubc13-Ub were

pooled and concentrated using Amicon or Pierce ultracentrifugation filter (3 kDa MWCO) and flash frozen in liquid nitrogen before storage at -80 °C.

**Synthesis of GGK**

Boc-GG (A152188) and Boc-Lys-tBu (A275334) were purchased from Ambeed. Boc-GG (422 mg, 1.82 mmol, 1.1 eq) and HATU (691 mg, 1.82 mmol, 1.1 eq) were dissolved in 50 mL tetrahydrofuran (THF) and DIPEA (1.5 mL, 9.1 mmol, 5.5 eq) before the addition of Boc-Lys-tBu (502 mg, 1.65 mmol, 1 eq). Reaction was stirred at room temperature overnight. Reaction completion was checked by TLC and mass spectrometry analysis, and volatiles were removed under reduced pressure. Solids were redissolved in ethyl acetate and washed 3 x with 0.5 M HCl, and volatiles were removed under reduced pressure. Global deprotection was performed using 1:1 trifluoroacetic acid (TFA) and dichloromethane (DCM) at room temperature for 2 hours before removing volatiles under reduced pressure. Final product was validated by LC-MS. LCMS (ESI): Calculated for $C_{10}H_{21}N_4O_4$ $[M + H]^+$ = 261.2; observed 261.2.

**Synthesis of Biotin-Ahx-LPLTG and Biotin-Ahx-LPKTG**

Ahx-LPLTG peptide was synthesized at a 0.1 mmol scale via solid-phase peptide synthesis using CEM LibertyBlue 2.0 system on Rink Amide 2 MBHA resin. Peptides were synthesized using microwave assist with iterative cycles of deprotection in 20% piperidine in dimethylformamide (DMF), rinsing with DMF, and coupling with N,N-diisopropylcarbodiimide (DIC) and Ethyl cyanohydroxyiminoacetate (OxymaPure). Resin was N-terminally Fmoc deprotected before removal from the instrument. Peptide was biotinylated manually at the N-terminus by reacting Biotin (98 mg, 4eq) with HATU (152 mg, 4 eq) in 3 mL of 4% DIPEA in DMF. Peptide was deprotected and cleaved from the resin in 2.5% tri-isopropylsilane, 5% water, and 92.5% trifluoroacetic acid at room temperature for 1 h. Peptide was purified using reverse-phase chromatography on a C18 column via 5-95% gradient of acetonitrile with 0.1% TFA over 24 minutes. For use in assays, Biotin-Ahx-LPLTG was dissolved to 100 mM concentration in water before dilution to the final concentration used in the assay. Peptides were validated by LC-MS.

Calculated for Biotin-Ahx-LPLTG $C_{39}H_{66}N_8O_{10}S$ $[M + H]^+$ = 838.5; observed 838.5.

Calculated for Biotin-Ahx-LPKTG $C_{39}H_{67}N_9O_{10}S$ $[M + H]^+$ = 853.5; observed 853.5.

**Sortase-mediated conjugation of Biotin-Ahx-LPLTG to GGK peptide**

For peptide-based sortylation studies, GGK "acceptor" and Biotin-Ahx-LPLTG "donor" peptides were dissolved in water to a stock concentration of 100 mM. Sortase conjugation was conducted using a final concentration of 6 uM SrtA 7M, 60 uM GGK peptide, and 90 uM Biotin-Ahx-LPLTG peptide in mass-spectrometry friendly SrtA 7M buffer (150 mM NaCl, 50 mM sodium acetate, pH 7.5). Reactions were conducted at 37 °C for the indicated time and the reaction quenched with 10% v/v formic acid, testing to ensure pH < 4. Samples were centrifuged for 10 min x 21,130 rcf to pellet any precipitate prior to LCMS analysis. Analytical LC-MS was performed using a system comprised of an Agilent 1260 Infinity II HPLC instrument equipped with an Agilent InfinityLab LS/MSD XT MS detector with electrospray ionisation. The system ran with a positive and negative switching mode and UV diode array detector using an Agilent ZORBAX SB-C18 RRHT (50 mm × 4.6 mm × 1.8 µm) column and gradient elution with two binary solvent systems: MeCN/$H_2O$ or MeCN/$H_2O$ plus 0.1% formic acid. Mass spectra was analyzed using MestreNova software.

**Sortase-mediated conjugation of Biotin-Ahx-LPLTG to digested proteins**

Digested proteins were reacted at 50 uM in a mass-spec modified SrtA 7M buffer (150 mM NaCl, 50 mM ammonium acetate pH 7.5) with 120 uM Biotin-Ahx-LPLTG and 6 uM SrtA 7M at 37 °C overnight. Following conjugation, 1 ug of trypsin was added directly to the reaction mixture and incubated at 37 °C for 6 hours to overnight to digest residual SrtA 7M in the reaction prior to mass spectrometry analysis. Samples were centrifuged at 21100 rcf x 10 minutes and the soluble portion analyzed by mass spectrometry or used directly in the following pull-down experiment.

**Streptavidin pull-down of biotinylated peptides from *in vitro* purified proteins**

Streptavidin agarose resin (Thermo fisher) was equilibrated with 150 mM NaCl, 50 mM ammonium acetate pH 7.5. Biotinylated peptides from the previous reaction were diluted to 100 uL in the equilibration buffer and incubated with the resin at room temperature for a minimum of 1 hour. Flow through and 100 uL was with equilibration buffer was collected for analysis and before 3 sets of washes consisting of 1 x 100 uL 1 M NaCl, 1 x 100 uL 5 M NaCl, and 1 x 100 uL 0.1 M Glycine (pH 2) were also performed before the elution step. Biotinylated peptides were eluted using either elution buffer (8 M Gdn-HCl, 50 mM glycine pH 1.5) or non-invasively using SrtA 5M hydrolysis. For SrtA 5M cleavage, the resin was resuspended in 100 uL of a mass-spectrometry modified SrtA 5M buffer (150 mM NaCl, 50 mM ammonium acetate pH 7.5, 5 mM CaCl2) and SrtA 5M was added to the reaction mixture to a final concentration of 2 uM. The reaction mixture was incubated at 37 °C in a benchtop shaker overnight and eluted by centrifugation through a fritted column. Eluted fraction was subjected to a trypsin digest as above to remove residual SrtA 5M before mass spectrometry analysis. Samples were analyzed using an Agilent LC-QTOF 6546-XT equipped with an Agilent Peptide Plus column and data was processed using MassHunter Bioconfirm software.

**QTOF LC-MS/MS Analysis**

LC-MS/MS analyses were conducted on samples from recombinant proteins on an Agilent 1290 Infinity II UHPLC system coupled with an Agilent 6545XT AdvanceBio LC/Q-TOF system equipped with an Agilent Dual Jet Stream ESI source. LC separation was obtained with an Agilent AdvanceBio Peptide Mapping column (2.1 × 150 mm, 2.7 µm). Peptides were separated over a gradient of 1-95% acetonitrile in water (both containing 0.1% formic acid) over 45 minutes with a flow rate of 0.4 ml/min. MS and MS/MS data were acquired using the Q-TOF with the following parameters:

| | |
|---|---|
| Drying Gas | 11 L/min |
| Drying Gas Temperature | 325 °C |
| Sheath Gas Flow | 10 L/min |
| Sheath Gas Temperature | 325 °C |
| Nebulizer Pressure | 35 psi |
| Capillary Voltage | 4,000 V |
| Nozzle Voltage | 0 V |
| Fragmentor Voltage | 175 V |
| Acquisition Mode | AutoMS2 |

**Data Analysis**

Data analysis was performed using Agilent MassHunter BioConfirm. Mass spectrometry files were searched against the relevant protein sequences and the noted chemical adducts were included as variable modifications, as well as carbamidomethylation at cysteine and methionine oxidation. Trypsin was specified as the protease, allowing a maximum of two missed cleavages. A precursor mass tolerance of 5 ppm and a fragment mass tolerance of 20 ppm Da were used

**General mammalian cell culture techniques**

Proteomics samples were initially prepared using HEK293T whole cell lysates. Cells were cultured in DMEM supplemented with Penn/Strep and 10% fetal bovine serum (FBS). Prior to harvesting, cells were treated for 30 minutes with 5 µM each of MG-132 and PR-619. To harvest, media was removed and adherent cells were treated with 0.05% trypsin before collection via centrifugation and washed 2x with Dulbecco's phosphate-buffered saline (DPBS) prior to flash freezing and storage of the cell pellet at -80 °C until further use.

**Proteomics sample preparation**

Harvested cells were thawed on ice and lysed in a buffer consisting of 8 M urea, 75 mM NaCl, 50 mM TRIS pH 8.0, and 1 mM EDTA supplemented with protease inhibitor cocktail (Thermo). Cells were lysed on ice for 15 minutes followed by 5 seconds of vortexing, and this was repeated twice before sonication using 40% amplitude for 3 seconds on, 3 seconds off for 3 cycles. Lysates were cleared by centrifugation at 4 °C for 10 min x 20,000g. Supernatant containing soluble proteins was reduced with 10 mM DTT at 60 °C for 30 minutes. Proteins were then treated with 20 mM iodoacetamide at room temperature in the dark for 10 minutes. Reduced and alkylated proteins were buffer exchanged into 50 mM triethylammonium bicarbonate (TEABC) to reduce urea to <2 M final concentration. Proteins were digested overnight at 37 °C using trypsin in a 1:50 trypsin:lysate ratio.

Digested peptides were desalted using SepPak C18 desalting columns. Briefly, columns were equilibrated with 100% acetonitrile followed by water with 0.1% formic acid. Digested peptides were acidified to pH ≤ 3 before loading onto the column twice. Samples were washed twice with 0.1% formic acid in water before elution using 50% acetonitrile and 0.1% formic acid in water.

**Biotinylation and streptavidin pull-down of peptides from *cellular extracts***
Digested peptides were suspended in SrtA 7M buffer (150 mM NaCl, 50 mM TRIS pH 8.0) to a concentration of 1.5 mg/mL. Peptides were treated with 3 mM Biotin-Ahx-LPL/KTGG peptide in the presence of 50 uM SrtA 7M at 37 °C for 16 hours. Peptides were incubated for 2 hours at 4 °C with streptavidin agarose (Pierce) using 1 mL of equilibrated resin per mg of total peptides in solution. Following binding, flow through was collected and resin was washed 3x in PBS and 2x in MilliQ water.

For denaturing acid elution, resin was incubated at room temperature for 1 hour with 8 M guanidinium HCl, 50 mM glycine pH 1.5 prior to elution.

For sortase-mediated elution (using Biotin-Ahx-LPLTGG peptide), resin bound peptides were resuspended in SrtA 5M buffer (150 mM NaCl, 50 mM TRIS pH 8.0, 5 mM $CaCl_2$) and treated with 50 uM SrtA 5M at 37 °C for 16 hours.

For trypsin-mediated elution (using Biotin-Ahx-LPKTGG peptide), resin bound peptides were resuspended in 50 mM TEABC buffer and trypsin was added 1:50 to the total biotin peptides. Elution from peptides proceeded at 37 °C for 16 hours.

Following elution, solvents were removed from peptides and were stored at -20 °C prior to desalting. Peptides were resuspended in 0.1% formic acid in water (ensuring the pH ≤ 3) and desalted using C18 StageTips generated in-house. Briefly, C18 material was packed into 200 uL tips, activated with 100% acetonitrile, and equilibrated with 0.1% formic acid in water. Samples were loaded onto the C18 material twice prior to 2 washes* with 0.1% formic acid in water (*note that for Gdn HCl elutions, samples were subjected to 5 washes prior to elution). Samples were eluted using 50% acetonitrile with 0.1% formic acid in water. Eluted samples were dried under reduced pressure and stored at -20 °C prior to analysis.

**Orbitrap LC-MS/MS Data Acquisition**
The C18 cleaned peptides were analyzed on ThermoScientific Orbitrap Exploris 240 mass spectrometer interfaced with UltiMate 3000 HPLC and UHPLC Systems. Digested peptides were reconstituted in 0.1 % formic acid with a final concentration of 500 ng/µl and 1 µl of sample (500 ng) was loaded on the column. Peptides were separated on an analytical column (75 µm × 15 cm) at a flow rate of 300 nL/min using a gradient of 1–25 % solvent B (0.1 % formic acid in 100 % acetonitrile) for the first 100 minutes and 25–30 % for next 5 minutes, 30-70 % for 5 minutes, 70–1 % for next 5 minutes. The total run time was set to 120 min. The mass spectrometer was operated in data-dependent acquisition mode. A survey full scan MS (from m/z 400–1600) was acquired in the Orbitrap with a resolution of 6000 Normalized AGC target 300. Data were acquired in topN with 20 dependent scans. Fragmented used normalized collision energy of 37 % and was detected at a mass resolution of 1500. Dynamic exclusion was set for 8s with a 10 ppm mass window.

### Data Analysis:

#### Identification of Biotin-Ahx-LPLTGG-K Sites

To identify peptides carrying the +877.4732 Da adduct, DDA LC–MS/MS data were searched using FragPipe 23.1 with MSFragger 4.4.1 against the human UniProt proteome supplemented with common contaminants and reversed decoy sequences. Spectra were searched using a labile mass-offset strategy with mass offsets of 0 and +877.4732 Da. The +877.4732 Da mass shift was restricted to lysine residues and delta-mass localization was enabled. Trypsin specificity was used with up to three missed cleavages. Precursor and fragment mass tolerances were 20 ppm, isotope errors of 0, +1, and +2 were considered, and b- and y-type product ions were used for scoring. Carbamidomethylation of cysteine (+57.02146 Da) was specified as a fixed modification, with methionine oxidation (+15.9949 Da) and protein N-terminal acetylation (+42.0106 Da) included as variable modifications.

Because preliminary inspection of confidently assigned +877.4732 spectra indicated extensive loss of the intact adduct during HCD, the search was performed in MSFragger labile-search mode. A +114.04293 Da Gly-Gly fragment remainder was specified to allow b- and y-type product ions retaining a diGly remnant on the modified lysine to contribute to peptide scoring and site localization. Three characteristic low-mass diagnostic ions, m/z 340.168, 425.258, and 453.250, were also specified. Diagnostic-ion intensity was not used as a mandatory spectrum instead, diagnostic-ion abundance was evaluated independently after peptide-spectrum matching.

Peptide-spectrum matches were filtered to a PSM q-value $\leq$0.01. Modified lysine positions were mapped from the localized peptide position to the corresponding protein sequence position. Common contaminant assignments were removed before biological interpretation. Ubiquitination sites were cross-referenced by exact protein accession and lysine position against a reference human ubiquitinome dataset[6]. Sites absent from this reference dataset were additionally examined in public PTM resources and published ubiquitin-remnant datasets. Together, these analyses defined a high-confidence +877.4732 lysine-adduct dataset using three independent features: (i) the appropriate +877.4732 precursor mass shift localized to lysine, (ii) sequence-specific peptide-backbone fragmentation, supplemented by K+114.04293 remainder ions, and (iii) the characteristic 340.168/425.258/453.250 diagnostic-ion signature.

#### Identification of Biotin-Ahx-LPLTGG *N*-term Sites

To identify *N*-terminally ubiquitinated proteins, tandem mass spectra were searched using a custom sequence-encoded database strategy designed to represent the residual enrichment tag as a linear peptide extension. The human UniProt reference proteome (UP000005640) was modified by prepending LPLTGG to the *N* terminus of each protein sequence. For proteins beginning with methionine, both methionine-retained and, where appropriate, initiator-methionine-cleaved variants were included. Target-decoy competition was preserved by generating matched decoy entries in which the biological protein sequence was reversed while the invariant *N*-terminal LPLTGG sequence was retained.

Spectra were searched with MSFragger in FragPipe using strict tryptic specificity and up to two missed cleavages. The residual Biotin-Ahx moiety was specified as a variable protein N-terminal modification of +339.16165 Da, such that the complete encoded conjugate corresponded to Biotin-Ahx-LPLTGG–protein. Methionine oxidation (+15.9949 Da) and carbamidomethylation of cysteine (+57.02146 Da) were included as variable and fixed modifications, respectively. Searches used a ±20 ppm precursor window with mass calibration and optimization enabled; the optimized main search used a 10 ppm fragment tolerance, b/y ions, intensity transformation, and the top 200 fragment peaks per spectrum. Peptide-spectrum matches were rescored with Percolator and protein inference was performed with ProteinProphet/Philosopher using target-decoy filtering.

Candidate N-terminal ubiquitination events were required to contain the LPLTGG extension, the +339.16165 Da Biotin-Ahx modification at the protein N terminus, and to map to protein position 1. Modified PSMs were then manually inspected for both peptide-sequence fragmentation and tag-specific diagnostic ions as above.

**TGG-modified peptide database searching and ubiquitination site analysis**

LC–MS/MS data were searched using FragPipe/MSFragger with parameters optimized for detection of the TGG remnant generated following tryptic release of sortase-enriched ubiquitinated peptides. In this workflow, cleavage of the bifunctional Biotin-Ahx-LPKTG reagent leaves a TGG modification corresponding to a monoisotopic mass addition of +215.0906 Da at the original ubiquitination site. Searches were performed using strict trypsin specificity with two enzymatic termini, precursor charge states of 1–4, and a ±10 ppm precursor mass tolerance. The +215.0906-Da mass shift was restricted to lysine residues and peptide N-termini, consistent with detection of canonical lysine ubiquitination and potential N-terminal ubiquitination.

Because HCD fragmentation of TGG-modified peptides frequently resulted in loss of the intact TGG group, candidate spectra were evaluated primarily from precursor mass and unshifted peptide backbone fragmentation. During method development, searches incorporating +57.0215 and +114.0429-Da partial TGG remainder ions were evaluated independently; however, remainder ions were not required for site assignment because many confidently localized TGG peptides were supported predominantly by unshifted b/y ions. Candidate TGG PSMs were retained following the standard FragPipe filtering workflow and subsequently inspected at the site level. Repeated PSMs mapping to the same peptide and modified lysine were collapsed to a single ubiquitination-site identification, while the number of independent spectra and the highest Hyperscore were retained as supporting metrics. Identified sites were cross-referenced against the previously compiled human ubiquitination atlas[6]. Site-level assignments were classified using the atlas confidence designation (Gold, Silver, or unannotated).

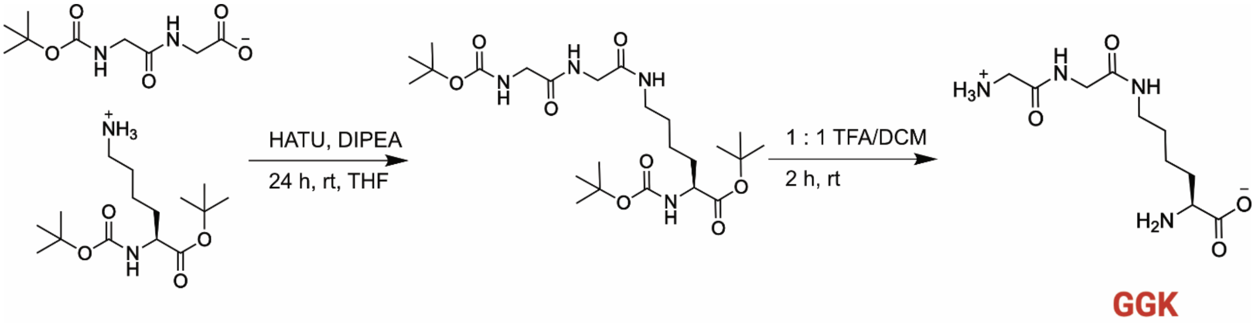


**Scheme S1:** Synthesis of GGK peptide.

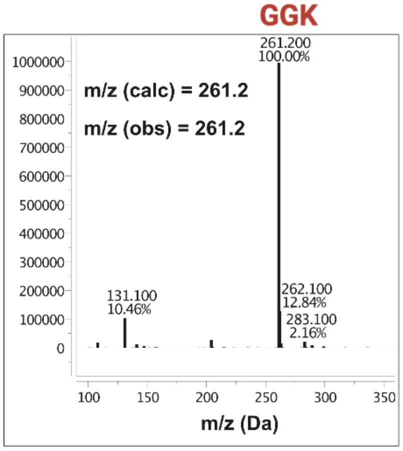


**Supplemental Figure 1:** LC-MS spectrum of GGK.

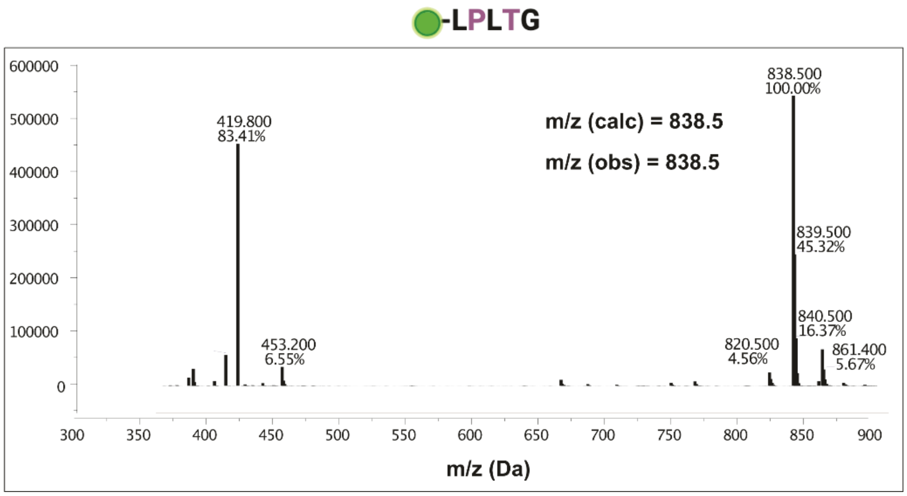


**Supplemental Figure 2:** LC-MS spectrum of Biotin-Ahx-LPLTG.

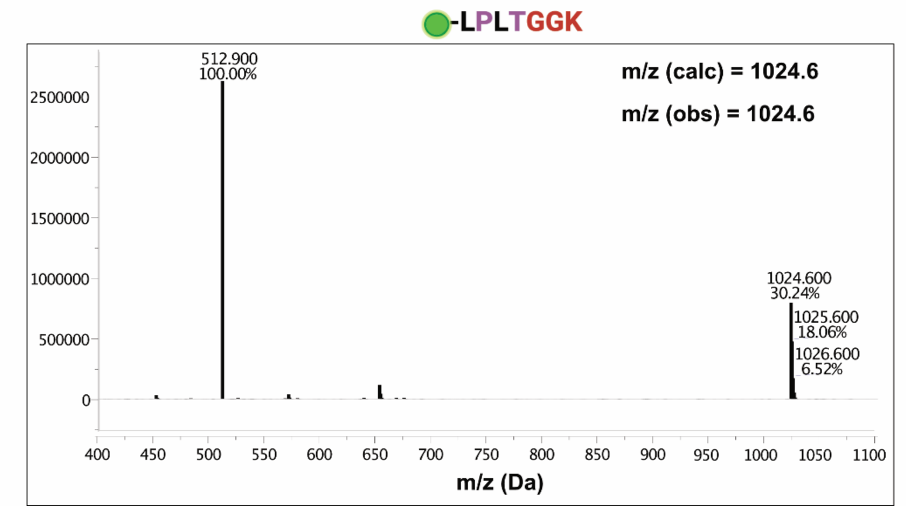


**Supplemental Figure 3:** LC-MS spectrum of Biotin-LPLTGGK product from Sortase reaction.

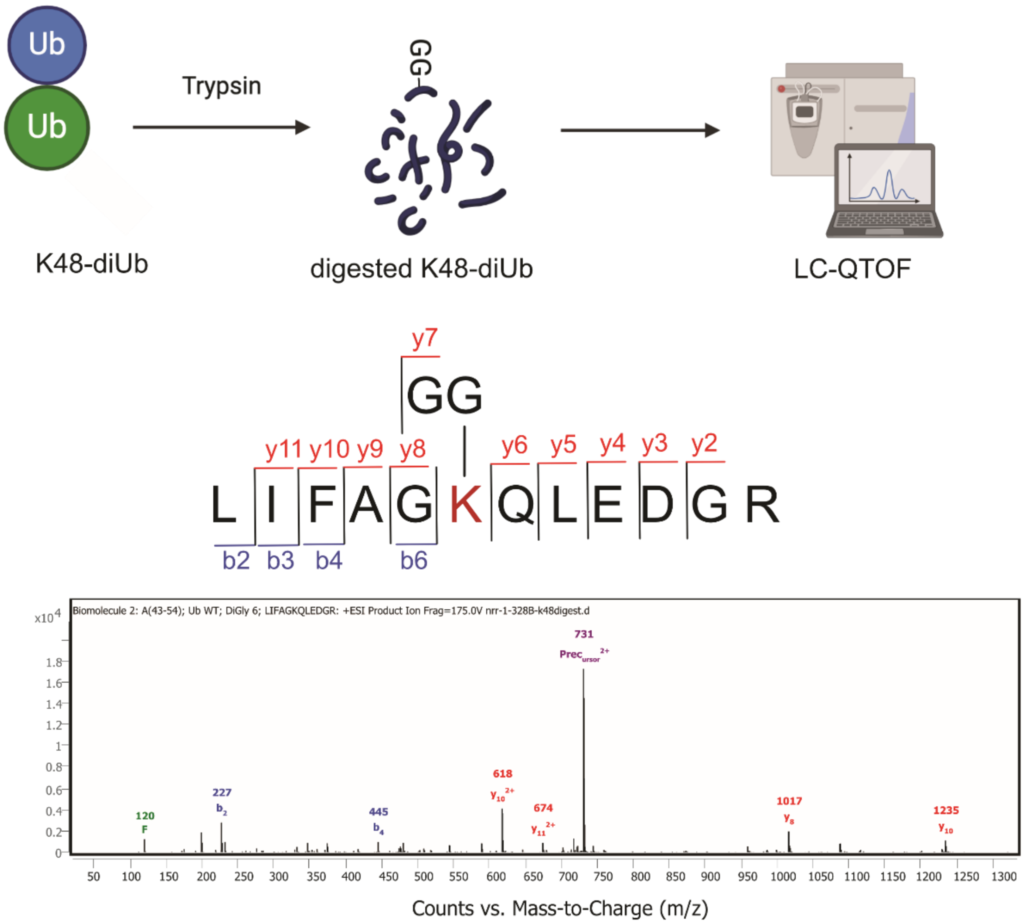


**Supplemental Figure 4:** Preparation, coverage, and LC-MS/MS spectrum of Ub K48 diglycyl remnant peptides.

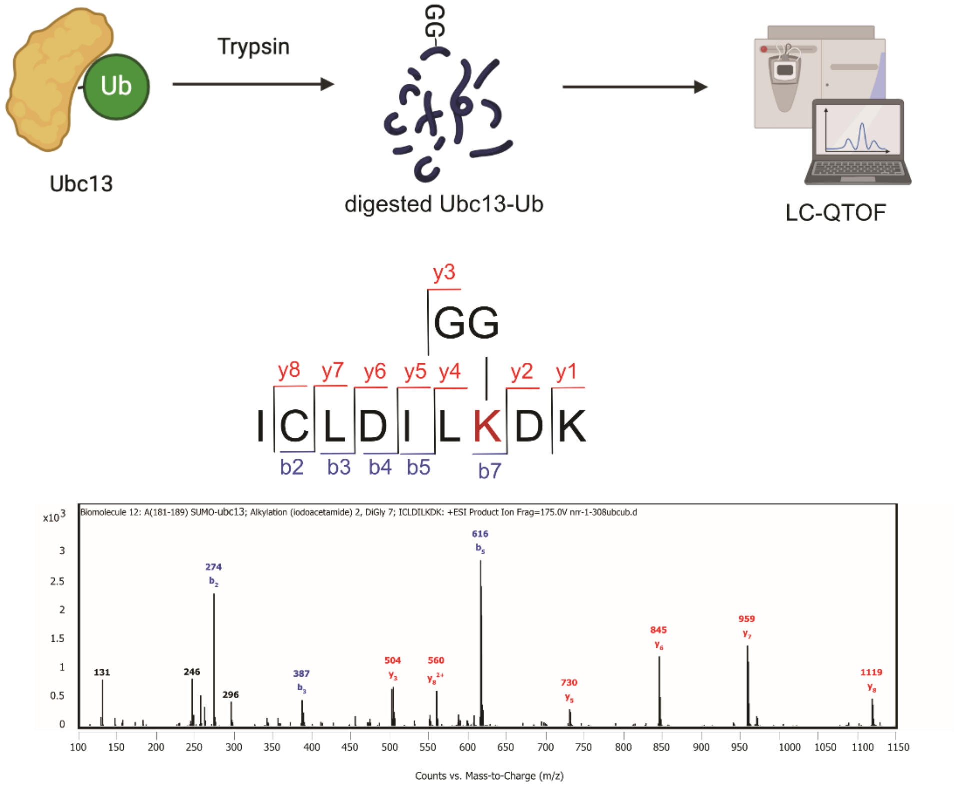


**Supplemental Figure 5:** Preparation, coverage, and LC-MS/MS spectrum of Ubc13 K92 diglycyl remnant peptides.
